# SUMOylation of the chromodomain factor MRG-1 in *C. elegans* affects chromatin-regulatory dynamics

**DOI:** 10.1101/2021.02.14.431134

**Authors:** Gülkiz Baytek, Alexander Blume, Funda Gerceker Demirel, Selman Bulut, Philipp Mertins, Baris Tursun

**Author notes:** Corresponding authors: Baris Tursun (BT), Gülkiz Baytek (GB).

## Abstract

Epigenetic mechanisms to control chromatin accessibility and structure is important for gene expression in eukaryotic cells. Chromatin regulation ensures proper development and cell fate specification but is also essential later in life. Modifications of histone proteins as an integral component of chromatin can promote either gene expression or repression, respectively. Proteins containing specific domains such as the chromodomain recognize mono-, di- or tri-methylated lysine residues on histone H3. The chromodomain protein MRG-1 in *Caenorhabditis elegans* is the ortholog of mammalian MRG15, which belongs to the <u>M</u>ORF4 <u>R</u>elated <u>G</u>ene (MRG) family in humans. In *C. elegans* MRG-1 predominantly binds methylated histone H3 lysine residues at position 36 (H3K36me3). MRG-1 is important during germline maturation and for safeguarding the germ cell identity. However, it lacks enzymatic activity and depends on protein-protein interaction to cooperate with other factors to regulate chromatin. To elucidate the variety of MRG-1 interaction partners we performed in-depth protein-protein interaction analysis using immunoprecipitations coupled with mass-spectrometry. Besides previously described and novel interactions with other proteins, we also detected a strong association with the Small Ubiquitin-like Modifier (SUMO). Since SUMO is known to be attached to proteins in order to modulate the target proteins activity we assessed whether MRG-1 is post-translationally modified by SUMOylation. Notably, we provide evidence that MRG-1 is indeed SUMOylated and that this post-translational modification influences the chromatin-binding profile of MRG-1 in the *C. elegans* genome. Our presented study hints towards an important role of SUMOylation in the context of epigenetic regulation via the chromodomain protein MRG-1, which may be a conserved phenomenon also in mammalian species.

## Introduction

In eukaryotes epigenetic regulation based on modifying chromatin is essential to ensure orchestrated gene expression. Chromatin regulation ensures proper development and differentiation of specific cell types. Modification of histone proteins, which are an integral component of eukaryotic chromatin, are key processes for the formation of active euchromatin or repressive heterochromatin states to promote either gene expression or repression, respectively. Modification by methylation of lysine residues on the N-terminal tails of histone H3 provide a binding site for chromatin-regulating proteins that act as ‘reader’ (Patel, 2016). Specific motifs such as the chromodomain enable proteins to bind to methylated histone H3, which was first identified in Polycomb (Pc) proteins and the Heterochromatin Protein 1 (HP1) (DasGupta et al., 2020). The evolutionarily conserved (Pc) proteins and (HP1) recognize mono-, di- or tri-methylated lysine residues on histone H3 (H3K27me3 and H3K9me3) and play an important role in maintaining repressed chromatin states. Many of the chromodomain-containing proteins are involved in chromatin-modifiers and remodeling complexes such as NuRD histone deacetylase complex and NuA4 histone acetyl transferase.

The chromo domain MRG15 belongs to <u>M</u>ORF4 related <u>g</u>ene (MRG) family in humans. The first identified MRG family protein in humans is the Mortality factor on chromosome 4 (MORF4) (Bertram et al., 1999). MORF4 induces replicative senescence in immortal cell lines and was therefore named “mortality factor”. Shortly after the discovery of MORF4, seven other MRG domain-containing genes were identified (Bertram and Pereira-Smith, 2001). Only two out of seven genes, MORF4 related gene on chromosome 15 (MRG15) and X (MRGX), are transcribed while others have pseudogene characteristics. Except for the differences in the N-termini, the amino acid sequences of MRG15 and MRGX are highly similar to MORF4. The differential N-termini of the three MRG family transcripts is likely to be the reason for the different functions of these family members. MRG15 is composed of a chromodomain in its N-terminus followed by a nuclear localization signal and the MRG domain at the C terminus composed of five alpha-helices (Xie et al., 2015). In contrast, MORF4 is composed of a NLS and MRG domain, lacking the chromodomain. Previous similarity searches using the human MRG15 amino acid sequence based on the databases have revealed that the MRG domain is conserved across species. The MRG15 homolog Msl-3 (male-specific lethal 3) in *Drosophila* is important for dosage compensation during sex specification (Chen et al., 2010). In *C. elegans* MRG-1 is orthologous to MRG15 also with respect to their function in the germline as MRG-1 is implicated in maintaining the germline by regulating proliferation and safeguarding the germ cell identity (Dombecki et al., 2011; Gupta et al., 2015; Hajduskova et al., 2019; Takasaki et al., 2007). MRG-1 participates in the chromatin regulation by interacting with the histone methyltransferases, and HDA-1 and SIN-3 histone deacetylase complexes (Beurton et al., 2019; Hajduskova et al., 2019). MRG-1has been shown to bind to distinct chromatin marks with a strong preference for H3K36me3 and H3K4me3 (Hajduskova et al., 2019). Recently, MRG-1 was identified as an important regulator of genomic integrity by partitioning genome into subnuclear compartments (Cabianca et al., 2019).

Besides post-translational modifications (PTM) of histone proteins also other PTMs play a key role in chromatin regulation. Modification by the small ubiquitin-related modifier (SUMO) known as SUMOylation has been implicated in several chromatin regulation processes (Cubeñas-Potts and Matunis, 2013; Vartiainen, 2010). Transcription factors (TFs) and chromatin regulators, especially those that have functions in DNA replication, chromatin silencing, and the DNA damage response, comprise the most frequently SUMOylated class (Hendriks and Vertegaal, 2016). SUMOylation is a reversible process which is highly similar to ubiquitination in terms of the enzymes that are utilized. These enzymes consist of an E1 (AOS-1/UBA-2), an E2 (UBC-9), several E3 ligases (GEI-17). Unlike vertebrates and some plants, *C. elegans* expresses only a single SUMO protein, SMO-1 (Surana et al., 2017). In mammals, SUMO-2, unlike SUMO-1, can assemble poly-SUMO chains on different substrates in the same protein. For SMO-1, such poly-SUMO chain formation has not been reported *in vivo*, although poly-SMO-1 chains were observed *in-vitro* (Surana et al., 2017). These poly-SUMO chains were implicated in recruiting other proteins to give rise to more complex interactions (Psakhye et al., 2019). Interestingly, SUMO is also implicated on cell identity in the context of TF-mediated reprogramming to pluripotency (Cossec et al., 2018). Further, SUMOylation of the TF Gatad2a, which is part of the nucleosome remodeling and deacetylase (NuRD) complex, inhibits reprogramming to iPSCs by blocking assembly and stability of NuRD (Mor et al., 2018).

Here we show that the chromodomain protein MRG-1 in *C. elegans* is SUMOylated. Interestingly, acute depletion of SUMO alters genome-wide distribution of MRG-1 by changing its preference for binding specific histone modifications. Overall, in *C. elegans* SUMO is implicated in chromatin regulation by altering chromatin dynamics of the chromodomain protein MRG-1.

## Results

### In-depth analysis of MRG-1 protein interaction network

Studying protein interactions can unravel the molecular mechanism behind unknown pathways. To interrogate the interactions of chromatin regulators including MRG-1 in *C. elegans*, we optimized co-immunoprecipitation protocols followed by mass spectrometry (CoIP-MS), which we previously applied to study chromatin-regulating proteins (Hajduskova et al., 2019; Kolundzic et al., 2018). Label-free quantification (LFQ) analysis is highly suitable for interaction proteomics protocols as specific interactions are detected only upon significant enrichment. Based on comparison studies between stable isotope metabolic labeling (SILAC) and label-free approaches, LFQ attained as accurate quantification as SILAC (Eberl et al., 2013). Yet, due to the higher variability in sample preparation three or more replicates are required (Cox et al., 2014). For that reason, we increased the number of biological replicates for studying the protein-protein interaction of MRG-1 and the control samples (Figure 1A).

**Figure 1:**
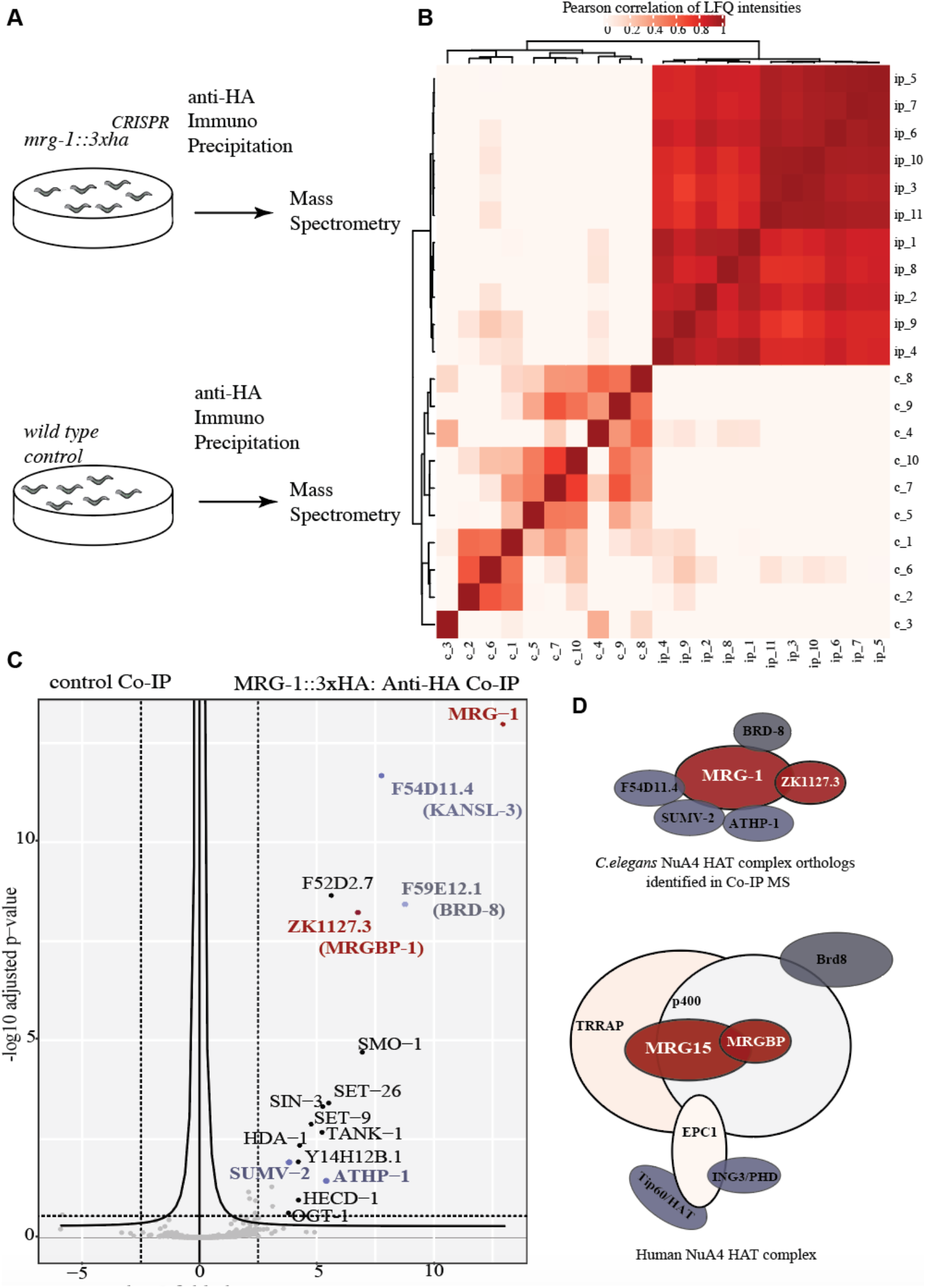
Protein-protein interaction studies revealed MRG-1’s interplay between NuA4 and HDAC complex. (A) Anti-HA co-immunoprecipitations (co-IPs) with subsequent LFQ mass spectrometry (IP-MS) to determine MRG-1 protein interactions. Wildtype (N2) with no HA tag was used as a negative control to eliminate background and *mrg-1::3xHA* (CRISPR/Cas9 generated) strain was used to identify MRG-1-interacting proteins. (B) Correlation matrix plotted as a heatmap, showing the Pearson correlations between 11 Co-IP and 10 control samples. (C) Volcano plot visualizing the enrichment of MRG-1 and its interaction partners between the IP and the control samples (x axis) and their corresponding adjusted p value (y axis). Statistics: t-test, adjusted P-value of 0.05 as false discovery rate cut-off. (D) Visualization of *C. elegans* NuA4 HAT complex orthologs identified in Co-IP MS and human NuA4 HAT complex (modified from Doyon Y. et al. 2004)

To get a confident set of interactors, we carried out 11 biological replicates for the IP condition and 10 replicates for the control (Figure 1A). A correlation matrix plotted as a heatmap, showing the Pearson correlations of LFQ intensities between 11 Co-IP and 10 control samples revealed more than 90% correlation among the IP samples (Figure 1B). A volcano plot demonstrates the enrichment of MRG-1 and its interaction partners compared to the control samples (x axis) (Figure 1C). The corresponding adjusted p values (y axis) were calculated by a *t-test* with 0.05 as false discovery rate cut-off (Figure 1C). Our results delivered MRG-1 interacting proteins, that were previously demonstrated as interaction partners of MRG-1 orthologues in other species (Figure 1D). As human MRG15 and yeast Eaf3 belong to Nucleosome Acetyltransferase of histone 4 (NuA4) (DasGupta et al., 2020), we checked whether *C. elegans* orthologs of NuA4 complex as MRG-1 interactors were enriched in our Co-IPs (Figure 1D). Notably, among the uncharacterized genes, F59E12.1 shows high homology to human bromodomain-containing protein BRD8, which is a component of NuA4 HAT complex (Cai et al., 2003). Another uncharacterized gene, ZK1127.3, expresses a protein that is homologous to human MRG-binding protein (MRGBP) and ZK1127.3’s deletion in *C. elegans* causes a larval lethal phenotype similar to *mrg-1* mutants (Barstead et al., 2012). Notably, F54D11.4 was one of the most significant interactors that is orthologous to human KAT8 regulatory NSL complex subunit 3 (KANSL3), which is encoded by the same gene family as SUMV-2 – another identified MRG-1 interactor (Figure 1C,D). Both F54D11.4 and SUMV-2 have histone acetyl transferase activity (HAT) that is necessary for the NuA4 complex (Cui et al., 2006). Moreover, ATHP-1, AT Hook plus PHD finger transcription factor may substitute for he PHD domain of the NuA4 complex in human. Overall, our CoIP-MS results concur with previous findings by revealing MRG-1-interactin proteins belonging to NuRD complex members, HDA-1 and SIN-3, as significant interactors of MRG-1 (Beurton et al., 2019).

### SUMOylation of MRG-1

Our in-depth CoIP-MS analysis revealed that SUMO, encoded by the gene *smo-1*, may interact with MRG-1. SUMO is attached to proteins as a post-translational modification that can cause important regulatory modulations of proteins (Celen and Sahin, 2020), we wondered whether MRG-1 may in fact be SUMOylated. To directly address whether MRG-1 is SUMOylated, we carried out Western Blot (WB) analysis with SMO-1 antibodies (DSHB Iowa) on immunoprecipitated MRG-1::3xHA (CRISPR/Cas9-tagged with 3xHA epitope). The presence of slower migrating MRG-1:3xHA proteins upon SMO-1 detection indicated that

MRG-1 is indeed SUMOylated (Figure 2A). Considering that one SMO-1 protein has a molecular weight of 11 kDa, the two bands corresponding to sumo-MRG-1 could be either an indication of two different SUMO sites or a multi-SUMOylation consisting of two SMO-1 proteins (Figure 2A).

**Figure 2:**
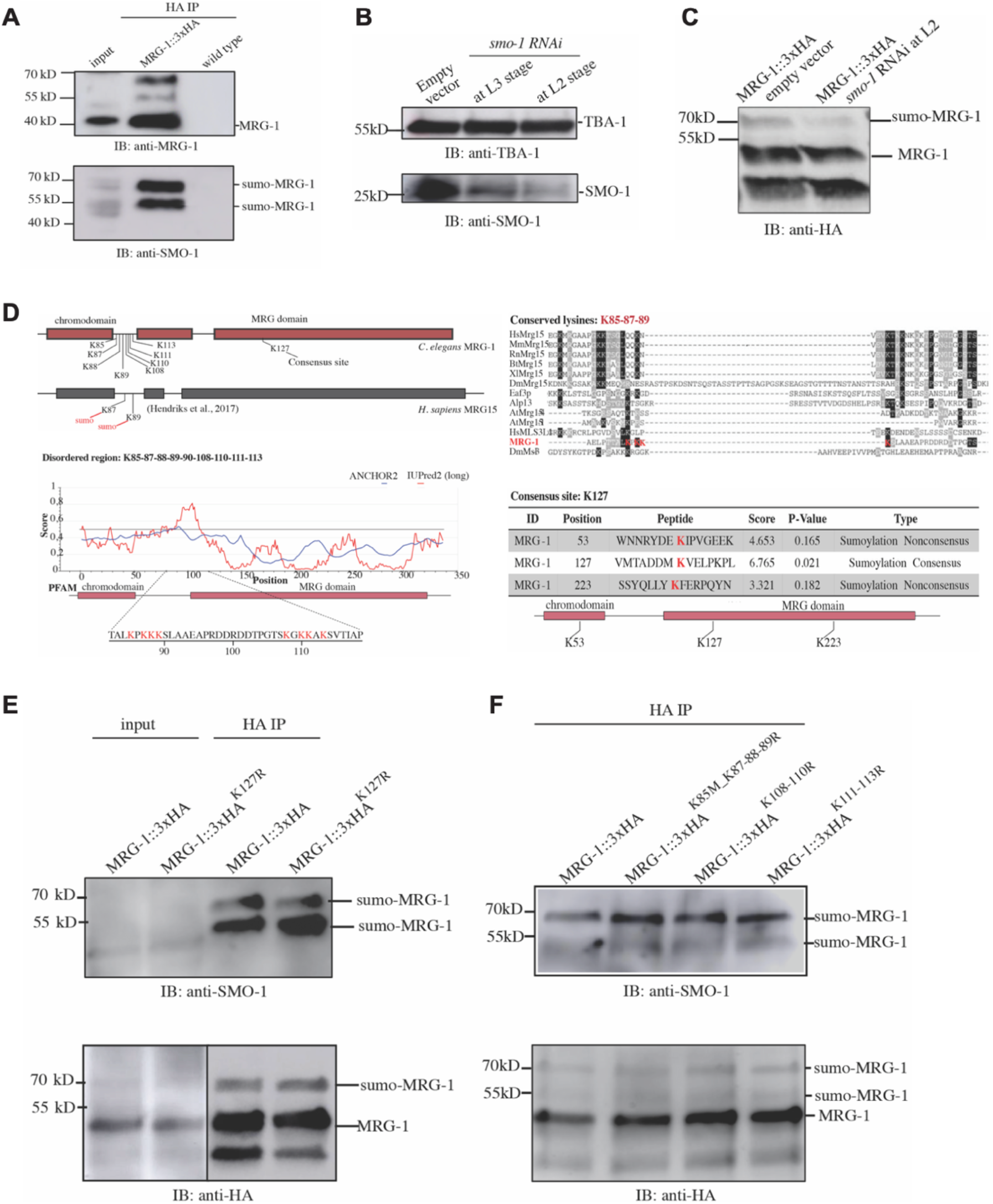
SUMOylation of MRG-1. (A) Immunoblotting of anti-HA IPs from *mrg-1::3xHA* and control strains with anti-MRG-1 and anti-SMO-1 antibodies, respectively, revealed SUMOylation of MRG-1. (B) Testing MRG-1 SUMOylation upon i RNAi. The efficiency of *smo-1* RNAi at different larval stages was tested to ensure proper development despite SUMO depletion. Tubulin was used as a loading control. Equal amounts of L2-staged *smo-1* RNAi exposed and control worms were picked at young adult stage and analyzed with anti-HA immunoblotting to see whether the corresponding sumo-MRG-1 band was downregulated. (C) Schematic representation of the comparison of predicted SUMO acceptor sites in the *C. elegans* MRG-1 and human MRG15. Based on Hendriks et al. 2017, 80% of sumo sites fall into disordered regions in human. K87 and K89 are the two SUMOylation sites on MRG15. In *C. elegans*, there is one candidate consensus SUMOylation site, K127. K85, K88, K108, K110, K111, and K113 residues were selected for CRISPR arginine mutations for being present in the disordered region. (D) Anti-HA Co-IP and the subsequent anti-HA and anti-SMO-1 western blot analysis of the mutated predicted SUMO lysines. CRISPR/Cas9-mediated lysine to arginine conversion of predicted SUMO-sites is not sufficient to deplete SUMOylation of MRG-1.

To provide further evidence that MRG-1 is SUMOylated, we tested whether global reduction of SMO-1 via RNAi would decrease MRG-1 SUMOylation (Figure 2B). First, the efficiency of the *smo-1* RNAi was tested. We took in account a previous report that *smo-1* F1 RNAi causes 100% embryonic lethality and that P0 RNAi at the L1 stage causes abrogated germline development and severe morphological aberrations (Jones et al., 2002). To bypass germline defect we exposed worms to *smo-1* RNAi at later stages of development such as L2 and L3 and worms were collected already at L4/young adult stage for WB analysis. The collected and lysed L4/young adult worms that were exposed to *smo-1* RNAi at the L2 stage expressed less SMO-1 protein than the worms exposed at the L3 stage (Figure 2B). L2 stage *smo-1* RNAi application caused some phenotypic aberrations related to tail morphology, while *smo-1* RNAi at the L3 stage caused no detectable developmental defects. Consequently, L2 *smo-1* RNAi significantly reduced the slower migrating MRG-1::3xHA bands in the WB analysis thereby providing further evidence that MRG-1 is posttranslationally modified by SUMO (Figure 2C). Next, we aimed to identify whether SUMOylation takes place at a specific lysine residue of MRG-1. We converted lysine residues of the MRG-1 protein into arginine via CRISPR/Cas9 gene editing (Figure 2C) to seek for a non-SUMOylated form of MRG-1. SUMOylated lysine residues are often embedded in the consensus sequence of ΨKxE (where Ψ is a hydrophobic residue and x any amino acid) (Celen and Sahin, 2020). We utilized a SUMOylation site prediction algorithm to target the lysine residues of MRG-1 which also considers that around 40% of SUMOylated sites are deviating from the consensus sequence (Zhao et al., 2014). Three sites of MRG-1 were predicted: K53, K127 and K223 - with K53 and K223 as non-consensus SUMOylation sites (Figure 2D). We decided to target K127 for arginine and additionally selected possible other target lysine residues that fall into disordered regions of MRG-1 protein because a previous study reported that the 81% enrichment of SUMO occurs in disordered regions (Hendriks et al., 2017). Interestingly, the lysine positions found in the disordered regions of MRG-1 were conserved and shown to be SUMOylated on human MRG15 (Figure 2D) (Bertram and Pereira-Smith, 2001; Hendriks et al., 2017).

All strains carrying CRISPR/Cas9-mediated mutations of *mrg-1* to convert lysine residues of MRG-1 protein we analyzed by anti-HA Co-IP with subsequent WB analysis (Figure 2E,F). The WB analysis with anti-SMO-1 antibody revealed that the mutated MRG-1::3xHA are still being SUMOylated *in vivo* (Figure 2E,F). This result indicated that either none of the mutated sites are the actual SUMOylation site, or that SUMOylation of MRG-1 may occur in a promiscuous manner concerning the MRG-1 lysine residues.

### SUMO depletion alters MRG-1 chromatin binding domains

Among 7,000 different SUMO targets identified to date, the transcription factors and chromatin regulators, in particular those that have functions in DNA replication, chromatin silencing, and the DNA damage response, comprise a frequently targeted class (Hendriks et al., 2017). Furthermore, a recent study showed that SUMO can function on chromatin as a ‘glue’ to maintain the cellular identity by stabilizing the epigenetic marks that are specific for somatic and pluripotent cells (Cossec et al., 2018).

The parallel functions in chromatin regulation of SUMO pathways and MRG-1 lead us to test whether SUMOylation has an impact MRG-1’s role as a chromatin factor. We performed anti-HA ChIP-seq using our MRG-1::3xHA strain with and without *smo-1* RNAi. As described before, we exposed animals at L2 stage to an acute *smo-1* RNAi for only 22 hours at 20°C till they reached the L4 stage (Figure 3A). As a control, animals treated with mock RNAi were used. Subsequent comparison of ChiP-seq data for MRG-1’s DNA-binding sites displayed significant differences upon acute exposure of *smo-1* RNAi from L2 to L4 stage as evidenced by Gene ontology (GO) analysis using gProfileR package (v0.7.0) (Raudvere et al., 2019) (Figure 3B). Chromatin sites that are less-bound by MRG-1 upon SUMO depletion are predominantly involved in cellular responses to external stimuli such as the temperature and unfolded protein responses (Figure 3B). Furthermore, genomic sites representing GO terms for histone deacetylases and nucleosomes, responses to DNA double strand breaks, and SIRT1 regulation of rRNA expression are also less bound by MRG-1 when SUMO is depleted for 22h (Figure 3B). For instance, histone genes such as *his-61* and *his-60* showed less MRG-1 occupancy suggesting that SUMOylation may coordinate MRG-1 binding to specific gene clusters (Figure 3C). Also, other specific genes encoding for E3 ubiquitin protein-ligase TRIM23 orthologs ZK1240.8 / ZK1240.1 and the *hsp-70* heat shock gene were less occupied by MRG-1 in response to *smo-1* RNAi reduction (Figure 3C).

**Figure 3:**
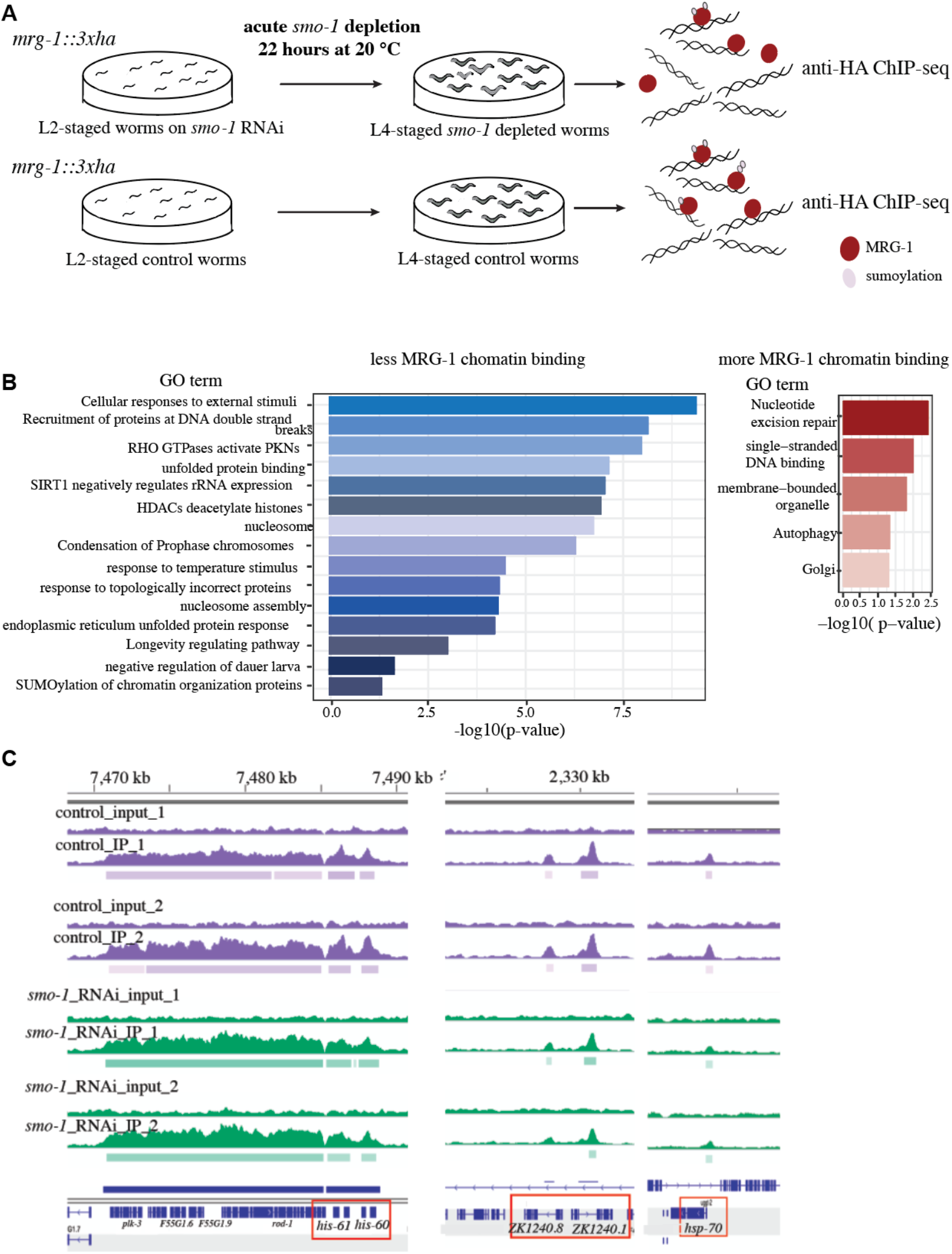
Effect of global SUMO depletion on the chromatin binding of MRG-1. (A) SUMO depletion was performed by *smo-1* RNAi-feeding beginning at the L2 larval stage of *mrg-1::3xHA* worms. For control, worms were fed with control RNAi bacteria. Worms were lysed at L4 stage and subjected to anti-HA ChIP-seq. (B) Gene ontology term analysis using g:Profiler showing differential genomic sites with less binding of MRG-1 upon *smo-1* RNAi. (C) Example of specific gene loci with altered MRG-1 binding upon depletion of SUMO.

Overall, lack of SUMOylation as a “glue” to bring together components of different chromatin regulator complexes may hinder complex formation thereby causing a reduction in MRG-1 binding to specific chromatin sites.

### SUMO RNAi alters MRG-1’s preference for chromatin sites with specific histone modifications

Previously, we showed that MRG-1 associates predominantly with histone modifications such as H3K36me3, H3K9ac and H3K4me3 (Hajduskova et al., 2019). As lack of SUMO may also alter the genomic distribution of MRG-1 depending on its preference for specific histione modification, we analyzed MRG-1 occupancy with respect to modENCODE histone modification data sets as we described previously (Hajduskova et al., 2019). As an internal control the meta-region profiles of MRG-1 DNA binding sites with no change in binding were used. Our analysis revealed less-bound and more-bound MRG-1 at chromatin domains with specific histone modifications after *smo-1* depletion (Figure 4A). Less MRG-1-bound sites carry mainly the repressive histone mark H3K27me3 but not the active mark H3K36me3, which is under wild-type conditions predominantly bound by MRG-1 (Hajduskova et al., 2019) (Figure 4A,B).

**Figure 4:**
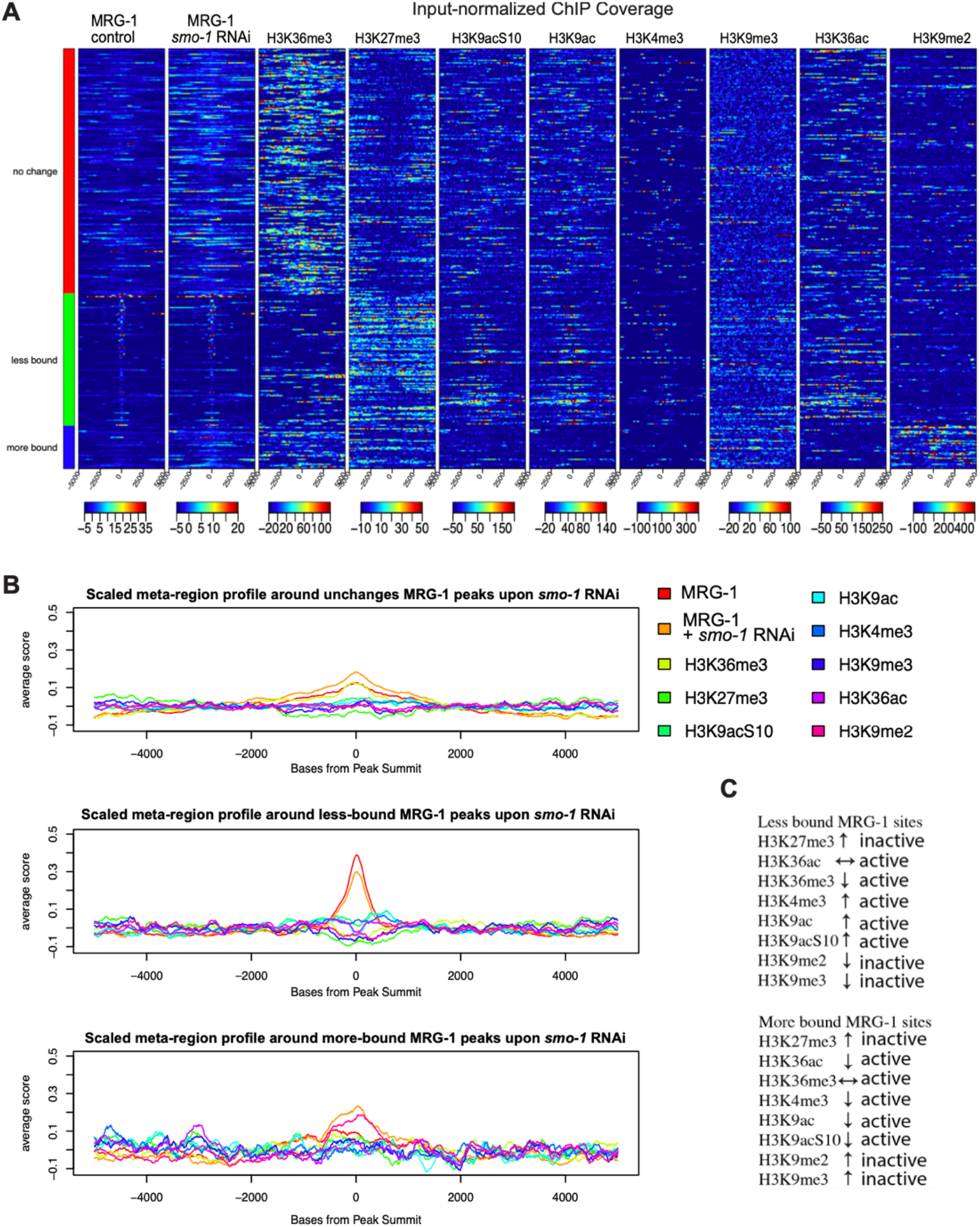
SUMO depletion alters MRG-1’s preference for chromatin sites with specific histone modifications. (A) Heatmaps show ChIP-seq signal for published histone modifications at 10kb windows centered around MRG-1 ChIP-seq peaks. The rows correspond to 200 randomly selected regions of unchanged peaks (red), 108 peaks showing less binding of MRG-1 upon *smo-1* RNAi (green) and 35 peaks out of the more bound (blue) peaks. Each color scale indicates the input-subtracted ChIP signal for respective matrix. Outlier values have been equalized to 30th and 99.7th percentiles for each matrix (B) he meta-region profiles for each of the three peak sets shows scaled and centered column-wise average signal over the respective peak regions plotted along the 10kb window. (C) Summary of MRG-1 binding changes upon SUMO depletion to histone modifications at differential loci compared to unchanged loci.

Overall, diminished SUMO in *C. elegans* leads to an altered chromatin-binding pattern of the chromodomain factor MRG-1. This alteration may be associated with different phenotypes observed upon SUMO depletion in *C. elegans* during development due to loss of the epigenome integrity.

## Discussion

Among the post-translation modifications, SUMOylation is a crucial mechanism sensing the external challenges that a cell can go through and regulates the function of its target proteins accordingly (Flotho and Melchior, 2013). There has been evidence showing global change in SUMOylation upon heat shock (Golebiowski et al., 2009; Miller et al., 2013), starvation (Shim et al., 2016), osmotic stress (Castro et al., 2012), DNA damage and hypoxia. Depending on their cellular function, while some of the SUMO target proteins are less SUMOylated, other are increasingly SUMOylated upon global stress. For example, proteins involved in chromatin remodeling complexes, mRNA splicing, and nucleosome were decreasingly SUMOylated in response to global stress (Hendriks et al., 2017). A large number of SUMO targets identified to date represent transcription factors (TFs), chromatin regulators, and DNA damage response factors, (Hendriks et al., 2017). Furthermore, SUMO is important to maintain cellular identities by stabilizing epigenetic marks that are specific for somatic and pluripotent cells (Cossec et al., 2018).

Our finding that MRG-1 in *C. elegans* is SUMOylated correspond with previous findings regarding the human homolog MRG15 (Hendriks et al., 2017). SUMOylation can act as a highly responsive pathway to external stimuli (Celen and Sahin, 2020) and may thereby coordinate MRG-1’s chromatin interaction. It is possible that chromatin binding of MRG-1 may be linked to sensing the external stimuli by SUMOylation in order to adjust gene expression. Interestingly, E3 ubiquitin protein-ligase TRIM23 orthologs ZK1240.8 and ZK1240.1 were both less occupied by MRG-1 in response to *smo-1* RNAi. Considering the direct interaction of MRG-1 with HECD-1 as revealed by our Co-IP results (Hajduskova et al., 2019) and the fact that MRG-1 is regulated by the ubiquitin-proteasome system (UPS) (Gupta et al., 2015) it I possible that SUMOylated MRG-1 is guided to specific chromatin sites to promote expression of proteasome-related genes in order to regulate its own protein levels. Such a negative feed-back loop could be important to respond to environmental changes sensed by SUMO or to prevent high levels of MRG-1 protein, which may disturb chromatin regulation.

Another interesting gene that had reduced MRG-1 binding upon *smo-1* RNAi is the *hsp-70* heat shock gene. The *hsp-70* gene encodes for a heat shock protein that is orthologous to human HSP70. Likewise, HSP70 is involved in the regulation of protein folding, degradation, responses to physiological and environmental stressors (Morimoto, 1998). Among genes that are more bound by MRG-1 upon *smo-1* RNAi are those that encode for regulators of nucleotide excision repair, single-stranded DNA binding, membrane-bounded organelle and autophagy. Giving the fact that MRG-1 is less bound to the histone and nucleosome-related genes upon SUMO depletion, MRG-1 may bind stronger to some genomic loci due to increased chromatin binding of different MRG-1-containing chromatin-regulation complexes. Moreover, reduced SUMOylation markedly changes MRG-1 dynamics on chromatin as it starts binding less to the sites with active histone mark H3K36me3, which in turn carry the repressive histone mark H3K27me3. Such changes in chromatin domain binding associated with specific repressive and active histone marks suggest that decreased SUMO levels may lead to decreased maintenance of the epigenetic signature of specific tissues. Moreover, while these effects can cause deregulated gene expression they could also increase the risk for chromatin defects and accumulation of chromosomal breaks as shown upon loss of MRG15 in human cells (Hayakawa et al., 2010). It is conceivable that MRG-1 has a similar role in maintaining genomic integrity, which has in fact been shown during meiosis in the *C. elegans* germline (Xu et al., 2012). As lack of SUMO in *C. elegans* leads to mitotic and meiotic defects (Reichman et al., 2018), it is likely that the SUMO pathway and MRG-1 converge with respect to maintaining genomic integrity in the germline.

## Material and Methods

### Worm strains

The wild-type *C. elegans* Bristol strain (N2) and CRISPR strains were maintained according to the standard protocol (Stiernagle 2006) at 20°C. BAT2012 *mrg-1(bar33[mrg-1*::*3xHA]) III*, (CRISPR/Cas9), BAT2126 *barSi36 [mrg-1_K127R:3xHA]* (CRISPR/Cas9), BAT2178 *barSi57 mrg-1K85R::3xHA* (CRISPR/Cas9), BAT2179 *barSi58 mrg-1K87R::3xHA* (CRISPR/Cas9), BAT2180 *barSi59 mrg-1K85M_K87-88-89R::3xHA* (CRISPR/Cas9), BAT2181 *barSi60 mrg-1K108_110R::3xHA* (CRISPR/Cas9), BAT2182 *barSi61 mrg-1K111_113R::3xHA* (CRISPR/Cas9).

### Synchronized worm population

Standard bleaching technique was applied to obtain synchronized worms. Gravid hermaphrodites were treated with sodium hypochlorite solution as previously described (Ahringer 2006). Household bleach (5% sodium hypochlorite) was mixed with 1 M NaOH and water in the 3:2:5 ratio. Worms were washed from NGM plates with M9 buffer, incubated in bleaching solution for 5 min in a 1:1 ratio and vortexed. Released embryos were washed three times with M9 buffer to completely remove hypochlorite. Embryos were incubated overnight to reach L1 stage. L1s were applied directly onto regular NGM plates for further maintenance of synchronized population.

### Western blot

Control and *smo-1* RNAi-treated worms were collected and washed with M9 buffer to remove bacteria. Samples were frozen at −20°C in SDS/PAGE sample buffer. Right before loading, samples were boiled for 10 min to denature the proteins and centrifuged for 10 min at full speed. MRG-1::3XHA was detected with rat anti-HA-HRP antibody (Roche) at a dilution of 1:500, SMO-1 was detected with anti-SUMO 6F2s mouse monoclonal from Hybridoma bank, and the secondary anti-mouse HRP antibody (catalog no. sc-2005; Santa Cruz Biotechnology) at 1:5.000 dilution.

As a standard loading control, we used anti-alpha-tubulin mouse monoclonal antibody from Sigma at 1:10000 dilution, and the secondary anti-mouse HRP antibody (catalog no. sc-2005; Santa Cruz Biotechnology) at 1:5.000 dilution. The Lumi Light detection kit (Roche) and the ImageQuant LAS4000 system (GE Healthcare Life Sciences) were used for the signal detection.

### Generation of CRISPR alleles

A PCR repair template introduced to worms by microinjection was used for CRISPR engineering containing the 3xHA tag sequence. Lysine to arginine mutations were inserted into the template along with a silent mutation for restriction site for convenient genotyping. The injection mix contained Cas9 protein (0.3 mg/μl), as well as a CRISPR RNA targeting mrg-1 (100 ng/μl). Overall, we used a recently described procedure (Dokshin et al., 2018). CRISPR RNA sequences are: GCAAGACTTTTCTTCTTCGG, ACGATCGTGATGACACTCCG, GAAATCGGTGACAATTGCGC.

### IP-MS

Each batch of biological replicates was performed in triplicate except the last with two which added up to 11 biological replicates. L4-stage wild-type and MRG-1::3xHA (CRISPR) worms were collected by M9 buffer, washed four times with M9 to get rid of bacteria, and concentrated into worm pellet after the last wash. The worms were added into liquid nitrogen drop by drop, by ensuring that the resulting “worm beads” did not exceed size of a black pepper to achieve even grinding afterwards. The frozen worms were then cryo-fractured using a pulverizer. To obtain a fine powder, worms were further ground using a mortar and pestle on dry ice. The worm powder was resuspended in 1.5× of lysis buffer (20 mM HEPES pH 7.4, 150 mM NaCl, 2 mM MgCl2, 0.1% Tween 20 and protease inhibitors), dounced with tight douncer 30 times and sonicated using a Biorupter (six times 30 sec ON, 30 sec OFF; high settings) followed by centrifugation at 16,000 × g at 4° for 10 min. The supernatant was removed to 2 ml Eppendorf tubes and incubated with HA antibodies (Roche) for 30 min on a rotator at 4°. Next, μMACS ProteinA beads (Miltenyi Biotec) were added into samples as instructed in the kit and samples were incubated for 30 min at 4° rotating. Meanwhile, the μMACS columns were placed to magnetic separator to be equilibrated and ready for sample application. Samples were diluted 5× of their volume with lysis buffer before being applied to columns and the columns with bound proteins were washed three times with lysis buffer to remove background binders. The proteins were eluted with elution buffer (100 mM Tris-Cl pH 6.8, 4% SDS, 20 mM DTT), heated to 95°. Eluted samples were prepared for mass spectrometry measurements by SP3 (Hughes et al. 2014), before analyzing on a Q Exactive Plus (Thermo Scientific) connected to a Proxeon HPLC system (Thermo Scientific). Label-free quantification was performed using MaxQuant as described below.

### IP-MS analysis

The raw mass spectrometry data were first analyzed using MaxQuant Software (Cox and Mann, 2008) and the resulting “proteinGroups.txt” was then processed using the Bioconductor R package DEP v1.0.1 following the section “Differential analysis” of the vignette (https://bioconductor.org/packages/release/bioc/vignettes/DEP/inst/doc/DEP.html#differential-analysis; version from November 17, 2017) with minor adjustments. First, we set the random seed to the number 123 to receive reproducible results and then we followed the paragraphs on “Loading of the Data” and “Data Preparation” of the vignette to create our raw protein table. Then we extracted the associated UniProt IDs from the raw protein table and queried them on the UniProt ID mapping tool (http://www.uniprot.org/uploadlists/) to generate a mapping from “UniProtKB AC/ID” to “Gene name,” which was downloaded as a mapping table in tsv format. The unmapped IDs were manually curated by a search in the UniProt Knowledgebase (UniProtKB) and then appended to the mapping table. This mapping table was loaded into R, where we first removed all rows of the table containing duplicated UniProt IDs. Next, we created unique gene names by appending to each duplicated gene name its number of occurrences separated by a dot, then we merged the raw protein table with the mapping table based on ID and UniProt ID, respectively, while keeping all rows of the raw protein table, and updated those entries in the names column where a gene name was available in the mapping table. Next, we loaded the table specifying the experimental design of the IP-MS analysis followed the instructions of the paragraph “Generate a SummarizedExperiment object” and “Filter on missing values,” where we decided to perform the less stringent filtering approach to keep those proteins that are identified in two out of three replicates of at least one condition. Then, we performed the steps described in “Normalization” and imputed the missing data using random draws from a manually defined left-shifted Gaussian distribution, with a shift of 1.8 and a scale of 0.3 as proposed in the “Impute data for missing values” paragraph. Next, we followed the paragraph “Differential enrichment analysis” to identify proteins being significantly enriched in comparison to the control co-immunoprecipitation with a minimum log2 fold change of 2 plus t-test with an adjusted P-value (α) < 0.05 (see Table S5).

### ChIP-seq

Not to intervene with MRG-1‘s function in the germline, we aimed to find an acute exposure where we can see the effects of *smo-1* RNAi on MRG-1’s chromatin binding. Instead of subjecting L1 worms to *smo-1* RNAi where the worms do not develop any germline, worms at later stages of development such as L2 and L3 were applied to *smo-1* RNAi to find a compromise between the depletion of SUMOylation and having intact worms with germline. For that, M9 arrested L1 worms were grown on OP50/Rluc plates until L2 or L3 stages at 20 °C. Then the L2 and L3 worms were washed off and transferred to smo-1 RNAi and control plates, respectively. L4/young adult animals were washed three times with M9 and fixed with 2% formaldehyde for 30 minutes followed by quenching with 0.125M glycine for 5 minutes. The samples were rinsed twice with PBS, and 100-200 ul of pellets were snap-frozen in liquid nitrogen and kept at −80°C. The pellets were washed once with 0.5 ml and resuspended in 1 ml FA Buffer (50nM HEPES/KOH pH 7.5, 1mM EDTA, 1% Triton X-100, 0.1 sodium deoxycholate, 150mM NaCl)+0.1% sarkosyl+protease inhibitor (Calbiochem) and then dounce-homogenized on ice with 30 strokes. The samples were sonicated using Bioruptor (Diagenode) with the setting of high power, 4°C, and 15 cycles, 30 sec on, 30 sec off. Soluble chromatin was isolated by centrifuging for 15 min at max speed at 4°C. The cellular debris was resuspended in 0.5 FA Buffer + 0.1% sarkosyl+protease inhibitor (Calbiochem) and sonicated again as described above. Isolated soluble chromatin was combined. The immunoprecipitation of HA-tag proteins was performed overnight in a total volume of 600 μl with 4 μg of HA-antibody (ab9110, Abcam) except the 5% of samples taken as input. Next day, 80 μl of Protein A Sepharose beads (Sigma) were added. The beads were washed with 1 ml of following buffers: twice with FA Buffer for 5 min, FA-1M NaCl for 5 min, FA-0.5M NaCl for 10 min, TEL Buffer (0,25M LiCL, 1% NP-40, 1% Sodium deoxycholate, 1mM EDTA, 10mM Tris-HCl pH 8.0) for 10 min, and twice with TE Buffer (pH8.0). DNA-protein complexes were eluted in 250 μl ChIP elution buffer (1%SDS, 250nM NaCl, 10 mM Tris pH8.0, 1mM EDTA) at 65°C for 30 min by shaking at 1400 rpm. The Inputs were treated for 5h with 20 μg of RNAse A (Invitrogen). The samples and inputs were treated with 10 μg of Proteinase K for 1h, and reverse cross-linked overnight at 65°C. DNA was purified with Qiagen MinElute PCR purification Kit (Qiagen).

The libraries were prepared using NEXTflex® qRNA-Seq™ Kit v2 (Bio Scientific) according to the manufacturer’s instructions. Libraries were sequenced using paired end sequencing length of 75 nucleotides on a HiSeq4000 machine (Illumina).

### ChIP-seq analysis

The ChIP-seq datasets for the *smo-1* RNAi comparisons have been analyzed using the PiGx-ChIPseq (Wurmus et al., 2018) pipeline (v0.0.40 and v0.0.51). Briefly, after trimming sequencing adapters with Trim-Galore (Krüger et al., 2015) reads have been mapped to version ce11 of the worm reference genome using bowtie2 (Langmead and Salzberg, 2012) ((v2.3.4.3, with ‘-k 1’ for smo-1-RNAi experiment) and a genome-wide coverage signal track was exported as BigWig file using a script based on Bioconductor R packages Rsamtools (Morgan et al. 2020), GenomicAlignments (Lawrence et al. 2013) and rtracklayer (Lawrence et al. 2009).

Subsequently reads were used for calling peaks for each sample with the MACS (version 2.1.1.20160309 and 2.2.6) (Zhang et al. 2008) submodule ‘callpeak’ with genome size set to the worm genome and automatic duplicate removal enabled for the *smo-1-RNAi* experiment, additionally skipping the model building process and extending the reads in 5′–3′ direction to 147 bp for the smo-1-RNAi experiment (‘-keep-dup auto -q 0.05’ and ‘-keep-dup auto -q 0.05 nomodel:” extsize: 147’ for smo-1-RNAi).

The number of reads mapping to the peaks for each sample were determined using the featureCounts function of the RsubRead (v2.0.1 and v2.4.2, for *smo-1* RNAi) (Liao et al. 2019) with default settings in single-end mode for the *smo-1* RNAi experiment (‘isPaired=false’) and ignoring multi-mapping reads.

Differential peak occupancy has been calculated using DESeq2 package (Love et al. 2014) (v1.26.0 and v1.30.0) where peaks with less than two supporting samples have been filtered out and peaks were classified as being differential if they showed an adjusted p-value below 0.1, with more bound peaks indicated by a log2 fold change value of greater than 0.5 (for the smo-1-RNAi experiment) and less bound peaks indicated by log2 fold change of less than 0.5 (for the smo-1-RNAi experiment). Each group of differential peaks was scanned for enriched GO terms and pathways using gProfileR package (v0.7.0) (Raudvere et al., 2019) and bar plot figures for enriched terms were generated using ggplot2 (v3.3.2) (Wilkinson, 2011).

Peak heatmaps and meta-region profiles of our MRG-1 ChIP-Seq and public histone modification ChIP-seq data were created as previously described in Hajduskova et al. 2019, with some adjustments. Briefly, public wiggle signal data for a selection of histone modifications that shared experimental conditions was fetched from the modEncode database (Gerstein et al., 2010) and then lifted over from their original genome assembly ce6 to ce11 and exported to BigWig format on the fly using CrossMap v0.3.8 (Zhao et al., 2013). The heatmaps were prepared and plotted using the Bioconductor R package genomation v1.10.0 (Akalin et al. 2015), where the rows of the underlying matrices correspond to peak regions correspond to the set of a) 200 randomly sampled peaks out of the not changing peaks, b) 158 peaks out of the less bound peaks and c) 60 peaks out of the more bound peaks. Each of the original peaks was resized to a length of 10 kb, while fixing the peak center and subsequently trimming bases reaching out of the chromosome boundaries, to define the input windows of a ScoreMatrixList with the MRG-1 and histone ChIP-Seq bigwig files as targets to calculated the average signal over 10bp for 1000 bins. To normalize the ChIP IP data by their respective input data, input coverage levels had to be scaled up towards the coverage levels of IP data. To achieve this, two ScoreMatrixLists were created for IP and input files separately and a scaling factor was calculated by estimating effective library size at the outer edges of peak regions (sum up coverage over bins 1 to 200 and 800 to 1000) for each samples IP and input data, then dividing IP’s effective library size by input’s effective library size. Then for each sample, the input data ScoreMatrix was multiplied by the scaling factor and subtracted from the IP data ScoreMatrix, producing the final ScoreMatrixList used for plotting the heatmap. [Refering to “10kb_1kBins_heatmap_reordered_commonScale_winsorized<30-97>.pdf””] The heatmap was created using the multiHeatMatrix() function, but enforcing a common scale and limiting the visualization to data within the [30,97] percentiles of the supplied matrices, by equalizing everything below the 30th / above the 97th percentile values to the values of the 30th / the 97th percentile values (‘common.scale=TRUE, winsorize=c(30,97)’).

The meta-region profiles for each of the peak sets described above were acquired by scaling and centering the ScoreMatrixList, subsetting each matrix into the three respective groups and then plotting per group for each sample the column-wise averages over the 10 bp bins.

### Genome Browser shot

The Genome Browser view shows the region “chr10:117,922,301-119,587,630” of the ce10 genome assembly and was created using the Bioconductor R package Gviz (Hahne and Ivanek, 2016).

## Acknowledgments

Thanks to Sergej Herzog for technical assistance and the CGC, funded by NIH P40OD010440, for providing strains. This work was partly sponsored by the ERC-StG-2014-637530 and the Max Delbrueck Center for Molecular Medicine in the Helmholtz Association. All procedures conducted in this study were approved by the Berlin State Department for Health and Social (LaGeSo). The authors declare that they have no competing interests.

